# Towards water-free biobanks: long-term dry-preservation at room temperature of desiccation-sensitive enzyme luciferase in air-dried insect cells

**DOI:** 10.1101/120063

**Authors:** Shingo Kikuta, Shunsuke J. Watanabe, Ryoichi Sato, Oleg Gusev, Alexander Nesmelov, Yoichiro Sogame, Richard Cornette, Takahiro Kikawada

## Abstract

Desiccation-tolerant cultured cells Pv11 derived from the anhydrobiotic *Polypedilum vanderplanki* embryo endure complete desiccation because of their ametabolic state and resume their metabolism after rehydration. These features led us to develop a novel dry preservation technology for enzymes as it was still unclear whether Pv11 cells preserved an exogenous enzyme in the dry state. This study shows that Pv11 cells protect an exogenous desiccation-sensitive enzyme, luciferase, preserving the enzymatic activity even after dry storage for 372 days at room temperature. A process including pre-incubation with trehalose, dehydration, storage, and rehydration allowed Pv11 (Pvll-Luc) cells stably expressing luciferase to survive desiccation and still emit luminescence caused by luciferase after rehydration. Luminescence produced by luciferase in Pvll-Luc cells after rehydration did not significantly decrease in presence of a translation inhibitor, showing that the activity did not derive from *de novo* enzyme synthesis following the resumption of cell metabolism. These findings indicate that the surviving Pv11 cells almost completely protect luciferase during desiccation. Lacking of the preincubation step resulted in the loss of luciferase activity after rehydration. We showed that preincubation with trehalose associated to induction of desiccation-tolerant related genes in Pv11 cells allowed effective *in vivo* preservation of enzymes in the dry state.

## Introduction

Enzymes, biological catalysts mainly represented by proteins, promote the decrease of activation energy of chemical reactions^1^. An enzyme binds to a substrate in the active site and releases products. Enzyme-substrate complex is formed to allow both components to interact with each other. This high substrate specificity is due to the precise three-dimensional structure of the enzyme^2^. Physicochemically, irreversible damage induced by exogenous stresses such as acidic pH, heating and repeated freezing and thawing results in the deactivation of the enzyme^3,4^. To prevent such damage, some enzymes are preserved in freezing conditions with cryoprotectants such as glycerol and trehalose. Glycerol maintains enzyme activity at low temperature by forming hydrogen bonds with water molecules^5,6^. Trehalose, a non-reducing disaccharide composed of two glucose molecules, acts like a chemical chaperone inhibiting protein aggregation and denaturation^7^. The specific physical characteristics of trehalose facilitate water replacement and eventual vitrification^8^. Water replacement hypothesis suggest that trehalose instead of water forms hydrogen bonds with the protein surface, resulting in the maintenance of protein conformation and integrity^9^. Vitrification means that trehalose forms in a glassy matrix during dehydration, which restricts protein and ion mobilizations^10^. Because of these properties, trehalose is involved in the stabilization of lipase in *Humicola lanuginosa* in freeze-dried state^11^.

Freeze-drying (lyophilization) is utilized to stabilize the enzyme structure and is applicable for long-term storage in every aspect of medical, pharmaceutical and food sciences. The principle behind this application is the removal of frozen water from materials through sublimation. To obtain freeze-dried products, the process is carried out typically as follows: pretreatment; concentrating products; freezing materials below triple point; drying with partial pressure allowing liquid water to be removed from the materials. The freeze-drying technique is characterized by a set of complex operations, including freezing, drying, evaporation, and precise temperature control to avoid denaturation^12^. To control the process of freeze-drying, high-energy and expensive equipment are required^13^. In this study, we proposed a novel preservation technology for enzymes in a dry state without chilling steps.

Some microorganisms and invertebrates massively accumulate anhydroprotectants to survive under drought conditions^14-17^. The sleeping chironomid *Polypedilum vanderplanki* inhabits in temporary rock pools in Africa. Its larvae can tolerate almost all complete desiccation during the dry season^18^. The larvae dehydrate slowly for 48 h to enter an ametabolic desiccation-tolerant state, namely anhydrobiosis^19^. In the process of anhydrobiosis induction, they accumulate biomolecules such as trehalose, highly hydrophilic proteins, antioxidants and heat-shock proteins, which allow the larvae to endure severe desiccation in a state of no metabolism^20,21^. These molecules contribute to preserve cells against the physicochemical damages due to oxidative stress such as DNA damage, protein degradation, and cell disruption^22,23^. The cultured cells Pvll, derived from *P. vanderplanki* embryo showed tolerance to almost complete desiccation as well as larvae^24,25^. Several anhydroprotectants are probably accumulated intrinsically in the cells during dehydration. Pv11 cells completely desiccated at less than 10% of relative humidity (RH) resume their metabolic activity immediately after rehydration. Therefore, essential proteins involved in the basic metabolism for the cell survival are preserved despite the almost complete dehydration. These cells may be used to preserve a protein of interest in the dry state without relying on expensive electric power supply.

We used DNA electroporation for Pv11 cells and obtained Pv11 (Pv11-KH) cells stably expressing AcGFPl^25^. Pv11-KH cells aid in the stabilization of the molecular structure of exogenous green fluorescent protein (GFP) during severe dehydration. The three-dimensional structure of GFP shows a beta-barrel composed of 11 mostly antiparallel beta strands that shield from solvents the chromophore in the central alpha helix responsible for an auto-catalyzed cyclization/oxidation chromophore maturation reaction^26^. Because of this intrinsic stability of GFP, it is not clear whether desiccation-tolerant Pv11 cells can also preserve the activity of an exogenously expressed enzyme. In yeasts dehydrated at 60% RH for 2 days, the activity of transfected exogenous luciferase, a desiccation-sensitive enzyme, markedly decreased to 10% compared to that observed prior to dehydration. Accumulated trehalose, due to dehydration in yeast cells, contributed slightly to the preservation of luciferase activity^27^. However, enzyme preservation using yeast is insufficient for long-term storage in a dry state at ambient temperature in terms of duration and RH conditions for dehydration. Desiccation tolerance in yeast does not only rely on anhydroprotectants but also on the cell wall creating effective protection for constitutive proteins. The cell wall reduces mechanical stress occurring through changes in osmotic pressure due to desiccation^28^. However, even a yeast equipped with such physical properties can barely protect an enzyme in the dry state for long-term storage. Since completely dried Pv11 cells were capable to survive up to 251 days at room temperature^24^, intrinsically expressed anhydroprotectants in this system may contribute preserve exogenous enzymes *in vivo*. The present study shows that Pv11 cells derived from a desiccation-tolerant insect can effectively protect luciferase against severe desiccation.

## Results

### Development of a protocol for stable expression of recombinant luciferase in Pv11 cells

Based on the method for establishing Pv11-KH cell line^25^, we generated a novel stable Pv11 cell line. The coding region for AcGFP1 in the expression vector was replaced with that of the luciferase gene and transfected into Pv11 cells. We successfully obtained the Emerald luciferase-expressing Pv11 cell line designated as Pv11-Luc. Bioluminescence caused by luciferase in living Pv11-Luc cells was observed using LV200 bioluminescence microscopy (Olympus, Osaka, Japan). Normal Pv11 cells did not show luminescence in the presence of D-luciferin (Fig. 1A), while luminescence of Pv11-Luc cells was observed (Fig. 1B). We defined the linear-correlation variables as coefficient of determination between the cell number and luminescence intensity caused by luciferase as exogenous expression efficiency (Fig. 1C). The original Pv11 cells did not show luminescence (Fig. 1C). We next analysed cell viability and luciferase activity of dry preserved Pv11-Luc cells after rehydration.

**Figure 1.**
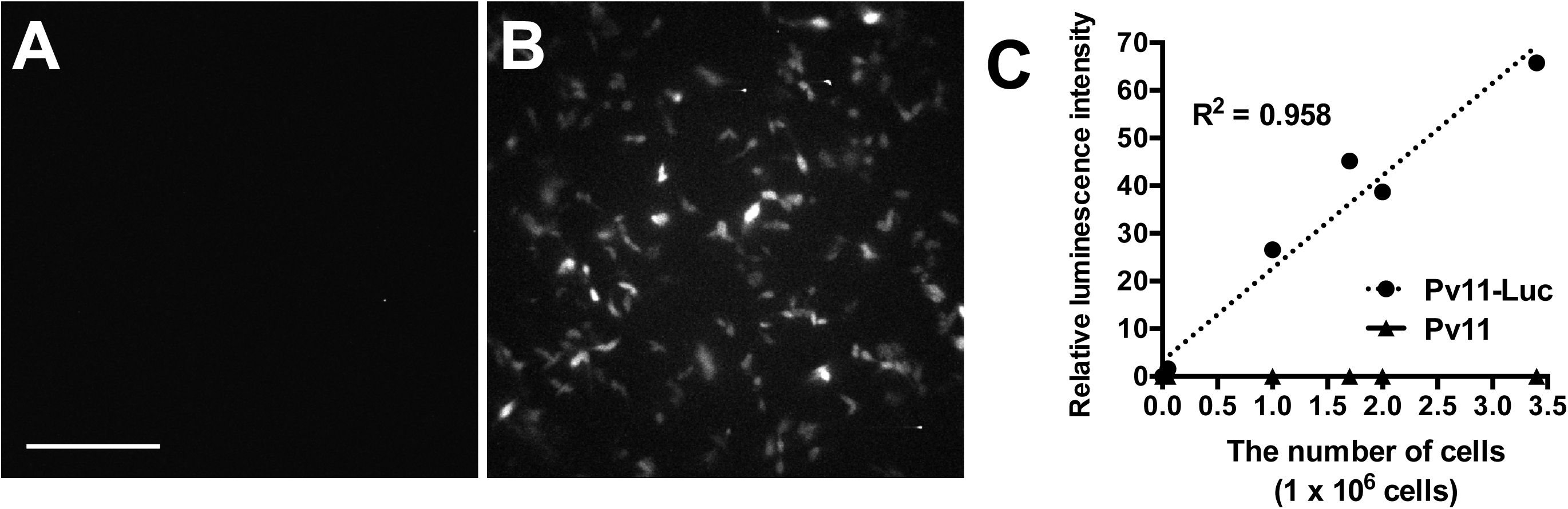
Generation of Pv11 (Pv11-Luc) cell line stably expressing luciferase. Pv11 cells transfected with the pPG121K-Luc plasmid were cultivated in insect medium with G418 for 3 weeks. Bioluminescence of normal Pv11 cells (**A**) and Pv11-Luc cells (**B**) was imaged in presence of 300 μΜ D-luciferin potassium salt. Exposure time was 4 min. Scale bar shows 100 μm. **C**. Luminescence intensity caused by luciferase was measured using Pv11-Luc cells and original Pv11 cells (i.e. absence of luciferase).

### Surviving Pv11-Luc cells retain luciferase activity even after severe dehydration

A suite of steps, including preincubation with trehalose, desiccation for 2 days, dry storage for 5 more days and subsequent rehydration was performed according to the schematic timeline (Fig. 2A). Luminescence intensity of rehydrated cells decreased compared to that of cells before dehydration, together with the reduction of cell viability (Fig. 2B). The reduction tendency of both cell viability and luciferase activity appeared to be well corresponding. The percentage of luciferase activity per surviving cell did not significantly different between before and after the desiccation/rehydration treatment (Fig. 2C; S2). The correlation between cell viability and luciferase activity was evaluated under various temperature conditions. We exposed desiccated Pv11-Luc cells to 25, 60 and 80°C for 5 min before rehydration, and then analysed cell viability and luciferase activity. The change of luciferase activity correlated strongly with the change of cell viability in response to temperatures (Fig. 2D). The results indicate that even after desiccation, surviving Pv11-Luc cells retain luciferase activity.

**Figure 2.**
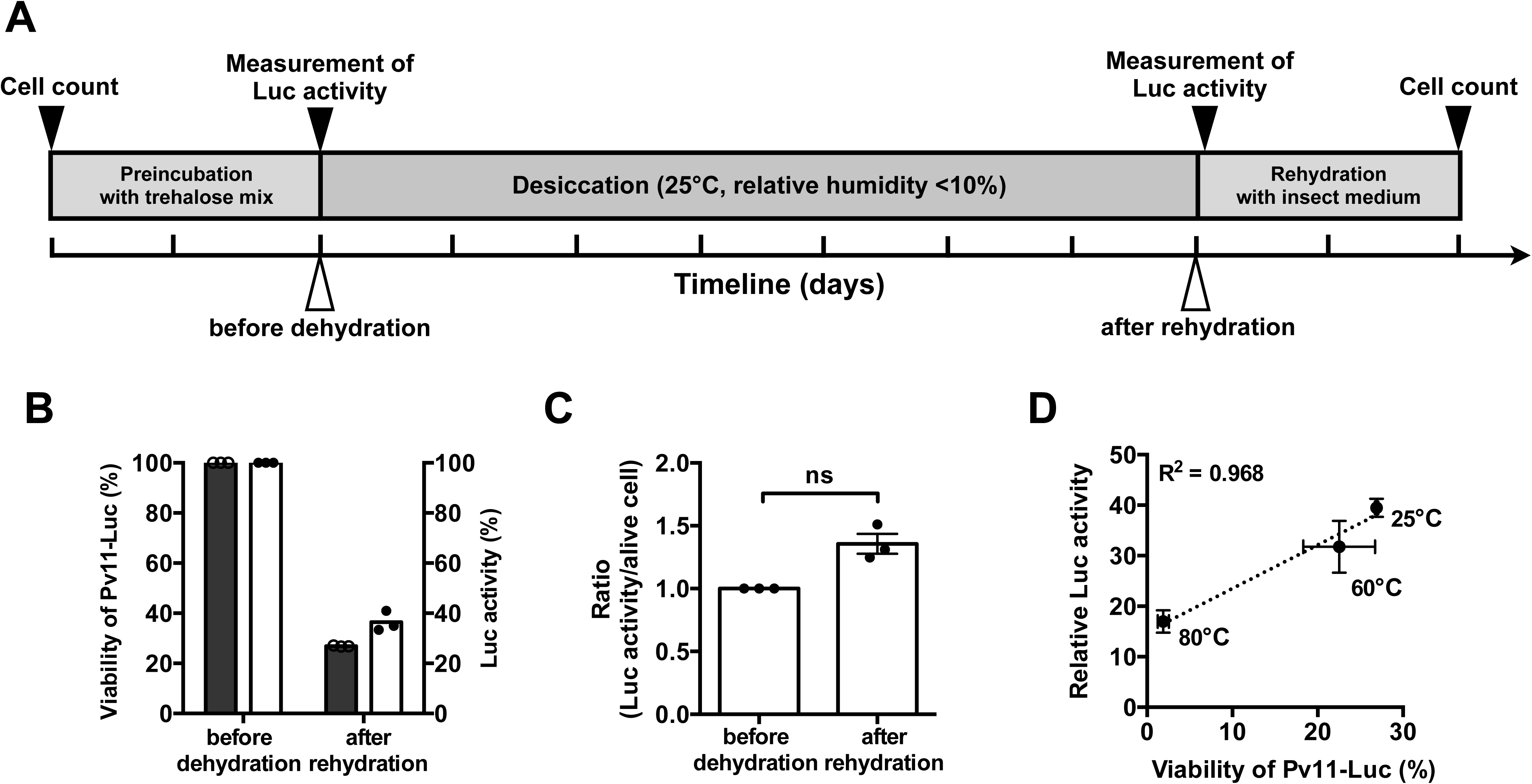
Viability and enzyme activity of Pv11 (Pv11-Luc) cells stably expressing luciferase. **A.** Schematic timeline represents the time point of each treatment (white arrowheads) and of the measurements (black arrowheads), respectively. Major tick intervals represent one day. The detailed procedures including preincubation, desiccation and rehydration are described in the previous study^24^. Pv11 cells are expected to be fully dehydrated about 30 h after the beginning of desiccation. Percentage of luminescence intensity of Luc and cell count was measured before the desiccation step (i.e., after pretreatment with 600 mM trehalose mixture containing both 10% IPL-41 and 1% FBS). Luminescence intensity caused by luciferase was measured 1 h after rehydration. Viability of Pv11-Luc cells was counted with the WST-8 assay. **B.** Viability of Pv11-Luc cells (left bar [black]) and percentage of Luc activity (right bar [white]) are shown as a scatter dot plot (n = 3). Error bar shows standard error of mean. **C.** Luc activity of surviving cells is represented as the ratio of luciferase activity over cell number. The number of surviving cells was estimated using the WST-8 colorimetric determination assay before dehydration and after rehydration. In this assay, “before dehydration” means before preincubation with trehalose mixture, and “after rehydration” means 2 days after rehydration with insect medium. The number of surviving cells was estimated using the calcein staining method after pre-incubation with trehalose and 1 h after rehydration (Fig. S2). Statistical significance was evaluated with Mann-Whitney *U* test (*p* = 0.064). D. Heat stress effect. Dry samples were heated for 5 min at 60 and 80°C before rehydration. The cells treated at 25°C represent unstressed control.

### Effect of long-term dry storage on luciferase activity

To induce desiccation tolerance in Pv11 cells, preincubation with the medium containing 600 mM trehalose before dehydration treatment is an indispensable step (Fig. 2A; Watanabe et al., 2016). Actually, the preincubation with trehalose allowed Pv11-Luc cells to survive desiccation for 7 days (Fig. 2A). Dry Pv11 cells can be preserved for 251 days maintaining cell viability^24^. We addressed whether Pv11-Luc cells retain luciferase activity even after long-term dry preservation. Desiccated Pv11-Luc cells kept for 372 days in the dry-state still showed luciferase activity although the survival rate dropped considerably (Fig. 3B).

**Figure 3.**
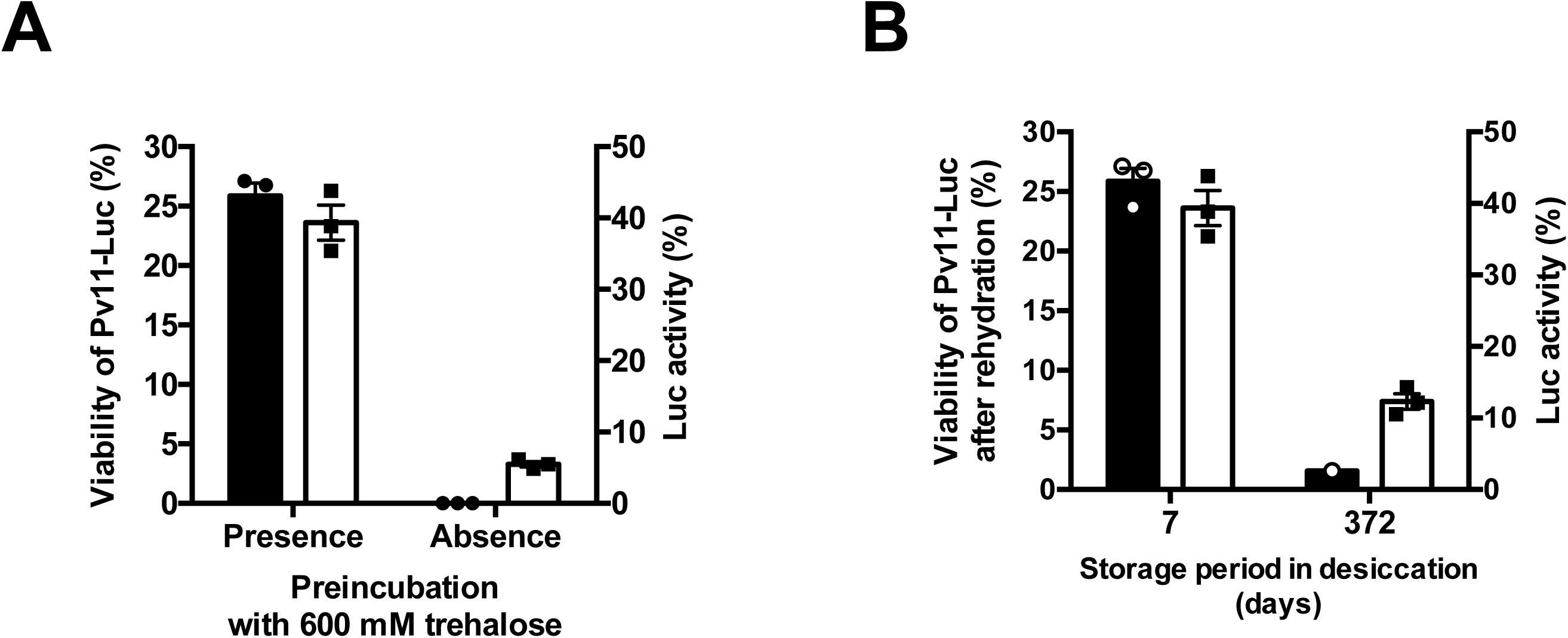
Effect of storage periods on Pv11-Luc cell survival after desiccation. Viability of dry-preserved Pv11-Luc cells (left bar [black]) and the corresponding relative Luc activity (right bar [white]) after rehydration are shown as a scatter dot plot (n = 3). Error bar shows standard error of mean. **A.** Effect of the preincubation with trehalose on cell viability and Luc activity. Desiccated Pv11-Luc cells that did or did not undergo trehalose pre-incubation were kept in a desiccator for 7 days. Cell viability and Luc activity were examined after rehydration, and are shown as a scatter dot plot. Error bar shows standard error of mean (n = 3). **B.** The long-term preservation of desiccated Pv11-Luc cells with trehalose pretreatment. Desiccated Pv11-Luc cells were kept for 7 and 372 days in a desiccator.

### Luciferase preservation in dry HEK293T cells

We examined whether a desiccation-sensitive mammalian HEK293T cells maintained luciferase activity after dry preservation (Fig. 4). The process of preincubation with trehalose, desiccation and rehydration was performed according to the same protocol used for Pv11-Luc cells. Luciferase expressing HEK293T cells did not show any indication of survival after dehydration (Fig. 4). Luminescence emitted by luciferase-expressing HEK293T cells was slightly observed after dehydration. The tendency of retaining luciferase activity during dehydration step was similar that observed for Pv11-Luc cells in the absence of preincubation with trehalose (Fig. 3A).

**Figure 4.**
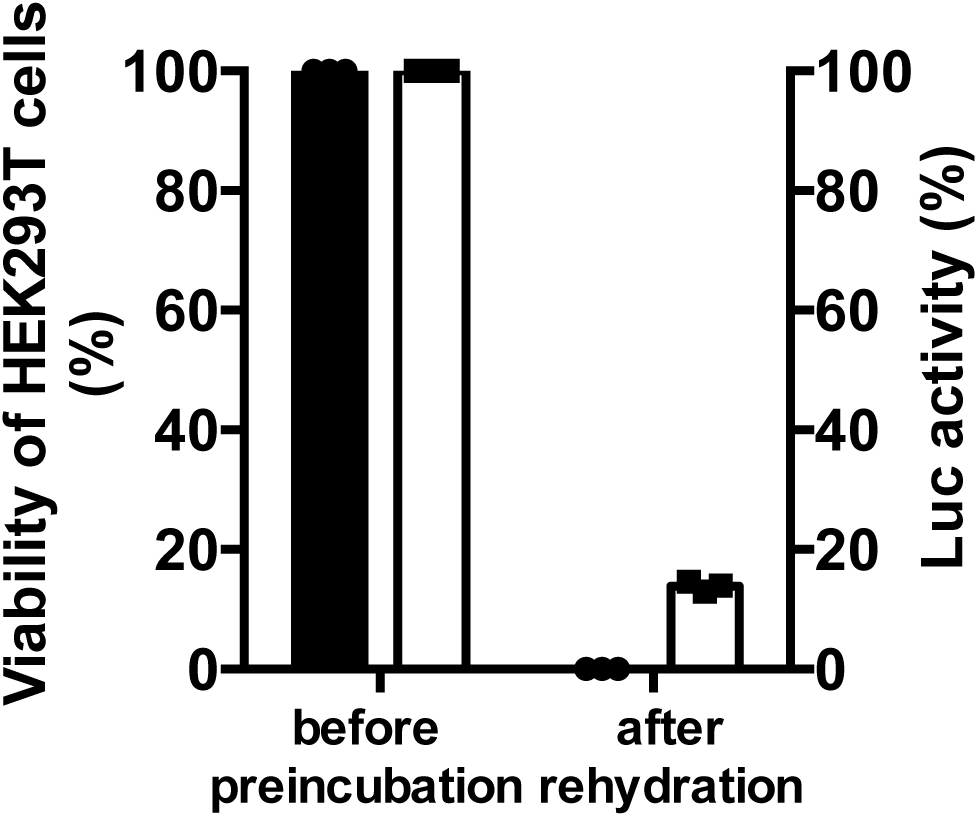
Cell viability and enzyme activity using a desiccation-sensitive mammalian cell line, HEK293T. HEK293T cells transiently expressing luciferase were preincubated with 600 mM trehalose and dehydrated following the same protocol used for Pv11 cells. Desiccated HEK293T cells were used for the measurement of cell viability and luciferase activity after rehydration. Viability of HEK293T-Luc cells (left bar [black]) and the corresponding relative Luc activity (right bar [white]) are shown as a scatter dot plot (n = 3).

### Recombinant luciferase preservation in the dry-state

To confirm the contribution of bovine serum albumin (BSA) and fetal bovine serum (FBS) to luciferase protection in the culture medium, we examined luciferase activity *in vitro* (Fig. 5). Recombinant luciferase was obtained using the *Escherichia coli* expression system (Fig. S1). Approximately 370 ng of luciferase in a 40-μL solution containing either BSA or FBS were dehydrated for 7 days following the same protocol for Pv11 cells. Luminescence was measured at 1 h after rehydration. The presence of 600 mM trehalose plus 10% IPL-41 insect medium (i.e. loss of both BSA and FBS) gave little effect of luciferase preservation. The presence of BSA at a molar ratio of 10:1 (BSA/Luc) also did not show significant enzyme protection compared to that observed in absence of BSA. Significant luciferase activity was observed in presence of BSA at a molar ratio of 50:1 (BSA/Luc). The presence of 10% FBS, equivalent of the concentration for cell dehydration, also showed luciferase activity of approximately 4% compared to that observed prior dehydration. In this *in vitro* assay, luciferase was partially preserved in BSA and FBS solutions. Both Pv11-Luc and HEK293T dead cells showed some luciferase activity after dry preservation (Fig 3A; 4); however, this luminescence caused by luciferase after dehydration was not because of protection by the cells themselves, but probably by BSA and FBS present in the medium.

**Figure 5.**
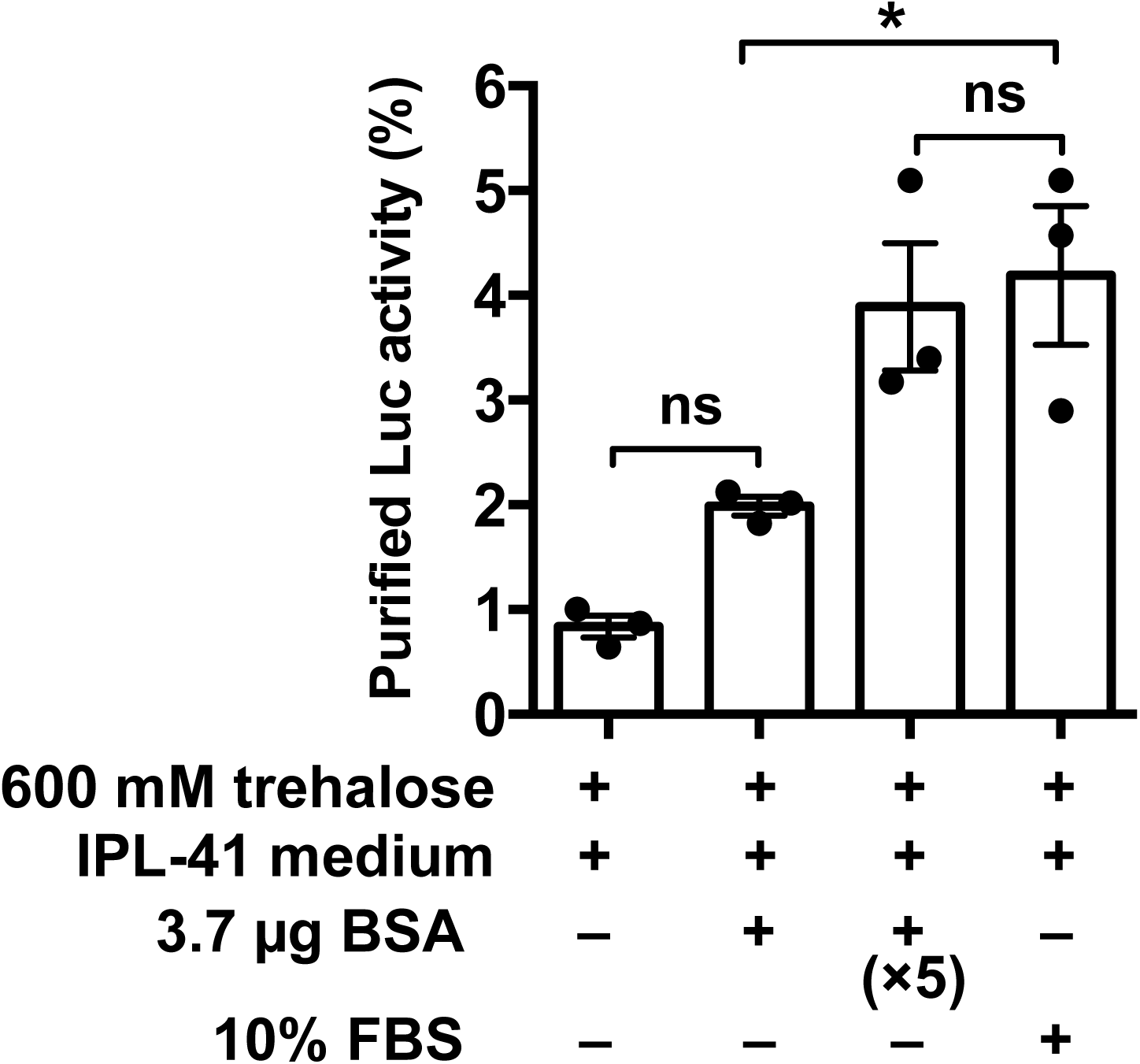
Enzyme activity of recombinant luciferase after dehydration. Emerald luciferase (370 ng) diluted into 40 μL of a 600mM trehalose solution containing either BSA, FBS or nothing was dehydrated for 7 days. The 3.7- μg BSA condition is represented as a molar ratio of 10:1 (BSA/Luc). Luciferase activity was measured after rehydration. Each value is shown as a scatter dot plot. Error bar shows standard error of mean (n = 3). Statistical significance was calculated by using one-way ANOVA and Tukey’s multiple comparison test as a post hoc analysis (“*” means significantly different *p* = 0.002; “ns” means not significantly different).

### Examination of luciferase preservation using a translation inhibitor

To examine whether luciferase activity was due to *de novo* synthesized enzyme after rehydration, we measured luminescence in the presence of a translation inhibitor. Both cycloheximide (CHX) at 0.35 mM and emetine at 1 mM were effectively inhibited protein synthesis (Fig. S3). Luminescence caused by luciferase in Pv11-Luc cells after rehydration did not significantly decrease in the presence of inhibitors compared to that observed for mock (Fig. 6). The results indicate that luciferase activity of Pv11-Luc cells after rehydration did not represent the activity of newly synthesized enzyme after resumption of cell metabolism. Altogether, the results show that surviving Pv11-Luc cells were able to preserve exogenously expressed luciferase in the dry state.

**Figure 6.**
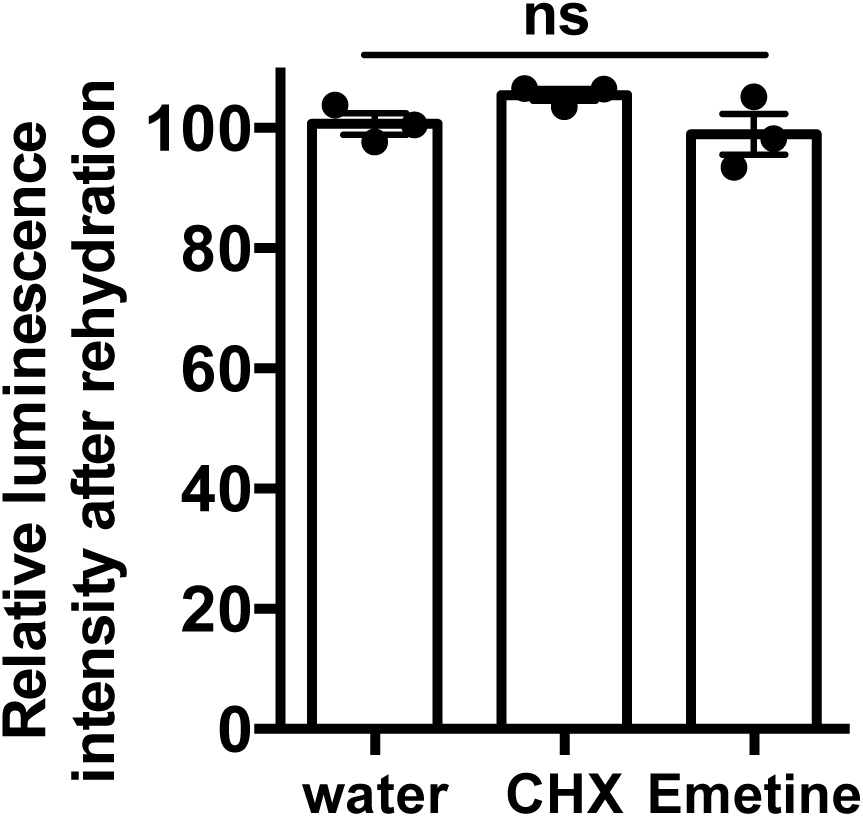
The *de novo* synthesis of luciferase after rehydration. Translation inhibitors cycloheximide (CHX, 0.35 mM) and emetine (1 mM), were added in the rehydration medium of dried Pv11-Luc cells. Luciferase activity was measured at 1 h after rehydration. Each value is shown as a scatter dot plot. Error bars show standard error of mean (n = 3). Statistical analyses were performed by using oneway ANOVA (*p* = 0.19). The effective concentration range of inhibitors for Pv11 cells was determined using the transient gene expression system (Fig. S3).

## Discussion

Stable luciferase expressing Pv11 were newly generated in order to examine the preservation of exogenous enzyme activity in the dry state. This study shows that desiccated Pv11-Luc cells protect the exogenously expressed luciferase against physicochemical damages occurring during drastic increase of endogenous ion concentrations and osmotic pressures due to desiccation stress. Pv11-Luc cells maintain the integrity of luciferase luciferase activity even after complete dehydration for long-term storage (Fig. 3B). The activity of luciferase after rehydration seems to be markedly decreased compared with that of the before dehydration (Fig. 2B). The decrease of luciferase activity could depend on the viability of Pv11-Luc cells during the process of dehydration/rehydration. Actually, the ratio showed that luciferase activity of surviving Pv11-Luc cells was not significantly different prior to dehydration and after rehydration (Fig. 2C; S2). Luciferase in Pv11 cells was not denatured and was preserved almost completely. Another explanation could be that the *de novo* synthesized luciferase occurring along with the resumption of cell metabolism after rehydration was responsible for the enzyme activity instead of the formerly protected enzyme. However, luminescence caused by luciferase in Pv11-Luc cells after rehydration did not significantly decrease in the presence of translation inhibitor compared to that observed for mock (Fig. 6), showing that the activity after rehydration did not represent the activity of luciferase generated from *de novo* synthesis. These findings suggest that Pv11-Luc cells that survived after exposure to desiccation stress protected luciferase, resulting in the detection of luciferase luminescence after rehydration.

Desiccation-intolerant mammalian cultured cells do not endogenously express genes encoding anhydroprotectants such as late embryogenesis-abundant (LEA) proteins, enzymes for synthesis of trehalose and trehalose transporter^21,29^. Even though the mammalian cells were pretreated with trehalose prior dehydration/rehydration as Pv11 cells, desiccation stress resulted in the loss of both cell viability and luciferase activity during desiccation (Fig. 4). Only a residual lusiferase activity was observed in HEK293T cells after rehydration in spite of the complete death of the cells. Similar result was also obtained with Pv11-Luc cells in absence of preincubation with trehalose (Fig. 4). We attempted to identify the factors responsible for the luciferase protection in dead cells. BSA, which is major component of FBS^30^ acts as a stabilizing agent for proteins^21^. Since the culture medium for both Pv11 and HEK293T contains FBS, we evaluated the contribution of BSA and FBS for luciferase protection *in vitro*. Bioluminescence caused by recombinant luciferase after dehydration drastically decreased to 1% compared to its original level in absence of BSA and FBS. Vitrification using IPL-41 insect medium and 600 mM trehalose gives little luciferase protection *in vitro* (Fig. 5). When both BSA and FBS were present in the medium, luciferase activity increased up to 4% compared to its original level. BSA and FBS contributed to protect luciferase against severe desiccation stress as demonstrated in previous *in vitro* assays^21^. Culture media containing FBS was used in all the steps from pre-incubation with trehalose to rehydration. Our results suggested that luminescence in dead cells did not derive from enzyme protection by BSA and FBS, or by similar protein-protein interactions within dead cells.

The enzyme drypreservation at 10% level for 2 days at 60% RH was accomplished in *Saccharomyces cerevisiae*, a eukaryote with desiccation tolerance^27^. In yeast, the cell wall has constitutive proteins potentially reducing the physical stress derived from the change in osmotic pressure during desiccation^28^. Although Pv11 cells derived from a eukaryotic organism showing desiccation tolerance similar to that of the yeast, they achieve anhydrobiosis without cell wall, and the mechanism of their desiccation tolerance is probably slightly different from that of the yeast^28,31^. Consequently, the cell wall acting structurally as a protection against osmotic pressure may not necessarily be required for the dry preservation of enzymes. Yeast and Pv11 cells require both trehalose and versatile chaperones as anhydroprotectants for their survival and enzyme preservation against desiccation^27^. Given that the surviving Pv11 cells allowed almost complete protection of exogenous luciferase in the dry state, cells should express very effective anhydroprotectants during dehydration. Enzyme preservation is achieved by the presence of intrinsic anhydroprotectants in Pv11 desiccation-tolerant insect cells. Anhydroprotectants, such as the chaperone-like LEA proteins acting as molecular shield, thioredoxin antioxidants, or small heat-shock proteins acting as anti-aggregants are expressed in *P. vanderplanki^32^*. Specifically, PvLEA4 contribute to inhibit abnormal protein aggregation of lactate dehydrogenase during desiccation *in vitro^21^*. These anhydroprotectants up-regulated under stress conditions^33^ allow enzyme preservation in desiccated cells. The biochemical role of each anhydroprotectant in Pv11 cells is not yet well understood. To date, the biochemical characteristics of PvLEA4 have been evaluated *in vitro* using recombinant proteins^21^. We believe that the measurement of luciferase activity in Pv11 cells will become a useful tool for studying anhydroprotectants in cells. For example, PvLEA3 is localized in the endoplasmic reticulum (ER)^34^. The ER targeting luciferase can be expressed in combination with the ER retention signal sequence. This approach should allow us to characterize the role of PvLEA3 in ER of Pv11 cells coupled with overexpression or clustered regularly interspaced short palindromic repeats (CRISPR)/Cas9 system.

Survival of *P. vanderplanki* larvae and Pv11 cells is accomplished through trehalose accumulation during desiccation. Gradual desiccation for 48 h allows trehalose to replace water and form a glassy matrix in the dry state. Vitrification of trehalose is inhibited over 65°C, which represents its glass transition temperature (*Tg*), resulting in the transformation from glassy to rubbery state and in the decrease of survival^8^. Cell viability and luciferase activity decreased at higher temperatures. This tendency was estimated by a correlation coefficient (Fig. 2D). Trehalose itself also acts as an anhydroprotectant of biomolecules in the glassy matrix and is indispensable for enzyme protection. Both trehalose and other anhydroprotectants are expected to act in combination to provide effective luciferase protection against desiccation stress in Pv11 cells.

Pvll cells immersed into 600 mM trehalose during dehydration formed a glassy matrix even without preincubation, however omission of the 600 mM trehalose preincubation step resulted in the loss of both cell viability and luciferase activity after rehydration (Fig. 3A). However, the physicochemical mechanism of trehalose implication in cell and enzyme protection has not been investigated. Preincubation with 600 mM trehalose in absence of 10% insect medium and 1% FBS did not allow cell survival after dehydration^24^. This suggested that the components in the medium and FBS, at least, are indispensable for Pv11 cells to endure the preincubation step with trehalose and exhibit desiccation tolerance. The preincubation step could have an alternative role, triggering the induction of anhydrobiosis-related genes in Pv11 cells. Given that the preincubation step allows Pv11 cells to protect the enzyme almost completely in the dry state, the preincubation mixture composed of 600 mM trehalose, 10% insect medium, and 1% FBS would faithfully up-regulate the anhydrobiosis-related genes for 48 h. Some anhydrobiotic genes such as *PvLea1, PvLea2* and *PvLea3* were up-regulated by salinity stress at 0.5% NaCl for 6 h^33^. Pv11 cells are conventionally cultivated using IPL-41 insect medium containing 0.3% NaCl^33,25^. Thus, the final salinity concentration is diluted to 0.03% in the preincubation mixture. Although the detailed mechanism of anhydrobiosis induction in Pv11 cells remains to be investigated, some stresses, such as salinity, oxidation, osmolation, or any combination of these may simulate Pv11 cells during the preincubation step and up-regulate anhydrobiosis-related genes. Understanding of this mechanism and improvement of the preincubation step should enhance the survival of Pv11 cells in the dry state and hopefully improve the protection of enzymes of interest for realistic production and dry storage in the future.

## Materials and methods

### Cultured cells

Desiccation-tolerant cell Pvll, an insect cultured cell derived from the sleeping chironomid *Polypedilum vanderplanki* embryo, was cultured with IPL-41 insect medium (Thermo Fisher Scientific, Carlsbad, CA), 10% fetal bovine serum (FBS, DS Pharma Biomedical Co., Ltd., Osaka, Japan), 2.6 g/L Bacto™ tryptose phosphate broth (BD, Franklin Lakes, NJ), 50 U/ml penicillin and 50 μg/ml streptomycin at 25°C. Human embryonic kidney 293T (HEK293T) cells were maintained with Dulbecco’s modified Eagle medium (D-MEM, Thermo Fisher Scientific), 10% fatal bovine serum (FBS, BioWest, Nuaillé, France), 4 mM GlutaMAX™ (Thermo Fisher Scientific), 100 U/ml penicillin and 100 μg/ml streptomycin under 5% CO2, 25°C, 95% humidity.

### Cell counting

The number of survived cells was measured using a counting chamber (Cell Science & Technology Inst. Inc., Miyagi, Japan) and the WST-8 colorimetric determination assay (Cell counting kit-8, Dojindo Molecular Technologies, Kumamoto, Japan). Optical density at 450 nm was measured using microplate-reader, iMark™ (Bio-Rad, Hercules, CA) at 25°C. To prevent color degradation, the measurements were examined in darkness. The WST-8 assay was performed according to the manufacturer's instructions. The surviving cell numbers were obtained from the standard curve of each experiment. Calcein-AM/Propidium Iodide double staining assays (Dojindo Molecular Technologies) for Pv11 cells were examined in accordance with the previous study^24^. The stained Pv11-Luc cells were observed using a fluorescence microscopy (BX53, Olympus, Osaka, Japan) equipped with a mercury lamp.

### Generation of luciferase expression construct for Pv11 cells

The coding regions of Emerald luciferase in pEluc-test (TOYOBO, Osaka, Japan) were obtained in the restriction enzyme treatments using *BgI*II and *XbaI*. The linear expression vector pPG121K of Pv11 cells was obtained in the restriction enzyme treatments using BamHI and XbaI. The PCR products of luciferase were directly introduced to the pPG121K vector with compatible restriction overhangs in *BgI*II and *Bam*HI. Sequence analyses for the confirmation of correctness of the constructs were performed by eurofins Genomics (Tokyo, Japan). Sequence data were analyzed with GENETYX-MAC ver. 16 software (GENETYX, Tokyo, Japan).

### Plasmid electroporation and luciferase stable Pv11 (Pv11-Luc) cell line

The pPG121K-Luc plasmid at 1~2 μg/μL was harvested using NucleoBond^®^ Xtra Midi EF (Macherey-Nagel, Düren, Germany). Approximately 10 μg plasmid DNA dissolved in 100 μL Opti-MEM^®^ (Thermo Fisher Scientific) was transfected into 1 × 10^7^ Pv11 cells in a cuvette with 2 mm gap (NepaGene, Chiba, Japan) using a NEPA21 super electroporator (NepaGene). The electroporation parameter was performed as follows: poring pulse 250 V, pulse length 4 msec, pulse interval 50 msec, number of pulse 6 times, positive discharge and 40% rate of decrease; transfer pulse 20V, pulse length 50 msec, pulse interval 50 msec, number of pulse 5 times, bipolar pulse and 40% rate of decrease. The Pv11 cells were cultured in a fresh IPL-41 insect medium containing FBS at 25°C after the electroporation. To generate luciferase stably-expressing Pv11 (Pv11-Luc), 400 μg/ml G418 (Thermo Fisher Scientific) was put into the cell culture at 2 days after the electroporation. The media containing G418 was exchanged every a week.

### Desiccation, anhydropreservation and rehydration procedures

The procedure of dry-preservation for Pv11 cells was described in the previous study^24,25^. In this study, the procedure was partially modified. 4 × 10^7^ cells were carried out for dehydration step. The cells were put into desiccation box (250 × 250 × 250 mm, AS ONE, UD-1, Japan) installed with 1 kg silica gel heated at 160°C for 3 h (Toyota Kako, Toyota Silica Gel, Japan), and kept at 25°C at less than 10% RH. The humidity inside the box was monitored using a humidity meter (TGK, Tokyo, Japan). Desiccated Pv11 cells were rehydrated with IPL-41 plus 10% FBS.

### Luciferase assay

Bioluminescence intensity by Emerald luciferase was acquired using a fluorescence/luminescence plate reader SH9000 (Hitachi High-Technologies Corp., Tokyo, Japan) at 25°C in a 96 well white microtiter plate (BD Falcon, Franklin Lakes, NJ). The integration time to obtain luminescence signals was set to 1.0 sec. Emerald Luc Luciferase Assay Reagent (TOYOBO) was used as manufacturer's instructions. Statistical analyses were conducted in Prism 6 (GraphPad, San Diego, CA). To obtain the luciferase activity of surviving Pv11-Luc cells after rehydration, Pv11 cells were collected in the centrifugation at 300 g for 5 min, and resuspended with PBS.

### Bioluminescence imaging of Pv11-Luc cells

Bioluminescence by luciferase was observed using Luminoview LV200 (Olympus, Osaka, Japan). Luminescence was acquired in MetaMorph^®^ imaging system (Molecular Devices, Sunnyvale, CA). Pv11 cells were attached onto a 35 mm glass bottom dish (AGC Techno Glass Co., Ltd., Tokyo, Japan) using 50 μg/ml cationic polymer, Polyethyleneimine “Max” MW 40000 (Polysciences, Inc., Warrington, PA). The cationic polymer was prepared as described previously^35^. D-luciferin potassium salt (Wako Pure Chemical Industries, Osaka, Japan) was dissolved in distilled water and kept at 4°C in the dark until use.

### Determination of concentration of translation inhibitors

Both cycloheximide (CHX, Sigma-Aldrich, St. Louis, MO) and emetine (Sigma-Aldrich) known as translation inhibitors for eukaryotic cells act by interfering elongation in protein synthesis. The concentration range of the translation inhibitors was referred to study^36^. To examine the effective concentration of the inhibitors for Pv11 cells, we confirmed the translation activity of Pv11 using the transient gene expression application. As a representative exogenous gene, AcGFP1 was used for the examinations. The pPGx2k-AcGFP1 expression vector was transfected into Pv11 cells using electroporation. The sequences and characteristics of the vector were shown in the previous study^25^. The electroporation parameter was set as referred above. The electroporated Pv11 cells were cultivated in the insect medium under various concentrations of inhibitor. We observed the AcGFP1 fluorescence at 48 h after the electroporation using a fluorescence microscopy (BX53, Olympus) and acquired the cytoplasmic fluorescence intensity in Metaview^®^ imaging system (Molecular Devices). Each inhibitor was inoculated into the rehydration buffer of Pv11-Luc cells. Luminescence by luciferase was measured at 1 h after rehydration.

### Purification of recombinant luciferase

To examine the preservation of purified Emerald luciferase in the dry-state, we expressed luciferase in *E. coli*. The coding regions of luciferase in pEluc-test (TOYOBO) were amplified by PCR primers containing restriction enzyme recognition sites as *Bam*HI and *Eco*RI, respectively using high-fidelity PCR polymerase PrimeSTAR^®^ HS (TaKaRa Bio). Forward primer was 5’– AGT<u>GGATCC</u>ATGGAGAGAGAGAAGAACGT–3’ and reverse primer was CTC<u>GAATTC</u>TTACAGCTTAGAAGCCTTCTC. Underlines represent restriction sites in the forward and reverse primer, respectively. After the restriction enzyme treatment, the PCR products were introduced into the bacterial expression vector pRSET-A (Thermo Fisher Scientific) applicable for the purification of 6xHis-tagged proteins. The pRSET-A-Luc plasmid was transferred into *E. coli* BL21 (DE3) (TaKaRa Bio). A single colony was picked and cultured in LB media containing 100 μg/ml ampicillin at 37°C until the optical density 600 (~0.6). 100 mM IPTG was put into the culture at 16°C, and kept for 6 h in darkness. The *E. coli* cells were disrupted by means of lysozyme treatment. Luciferase was purified using Ni–NTA chromatography, Profinity™-IMAC (Bio-Rad). Binding to resin in a column was washed with a buffer (20 mM Tris-HCl pH 8.0, 10 mM imidazole) and the recombinant luciferase was eluted with 20 mM Tris-HCl pH 8.0, 500 mM imidazole. Eluted proteins were concentrated with micro filtration (Merck Millipore, Darmstadt, Germany). The purity was confirmed by SDS-PAGE. The protein concentration was measured using Qubit^®^ Fluorometer (Thermo Fisher Scientific). Approximately 370 ng recombinant luciferase in 40 μL volume of trehalose mixture was used for desiccation, and it was kept for 7 days at 25°C in desiccation. The luminescence by luciferase was measured at 1 h after rehydration with PBS using the luminescence plate reader.

### Luciferase transient expression in HEK293T cells

We examined the preservation of luciferase activity in mammalian cells in a dry-state. The coding regions of luciferase in pEluc-test (TOYOBO) and the linear mammalian expression vector pcDNA3.1 (Thermo Fisher Scientific) were obtained in the restriction enzyme treatments using *EcoRV* and *XbaI*, respectively. The fragment encoded luciferase was introduced into the pcDNA3.1 vector. The Eluc expression vector was transfected into HEK293T cultured cells in accordance with the procedures in the previous study^37^. Desiccation of cells (4 × 10^7^ cells) was examined at 48 h after transfection. IPL-41 insect medium used for the preincubation with trehalose, dehydration and rehydration in each step was replaced with D-MEM for mammalian cell culture.

## Abbreviation

Luc: luciferase
GFP: green fluorescent protein
IPTG: isopropyl-β-D-thiogalactopyranoside
LEA: late embryogenesis abundant
HEK293T: human embryonic kidney 293T
BSA: bovine serum albumin
FBS: fetal bovine serum
D-MEM: Dulbecco's modified Eagle medium
ER: endoplasmic reticulum
RH: relative humidity

## Acknowledgment

This research was partially supported by Japan Society for the Promotion of Science KAKENHI (Grant Number 26850216 for SK; 16K15073 for TK) and by Russian Science Foundation grant for international groups (No. 14-44-00022). HEK293T cells were provided by RIKEN Cell Bank (Tsukuba, Japan). We thank Ms. Tomoe Shiratori (Institute of Agrobiological Sciences, NARO) for the preparation and maintenance of Pv11 cell lines.

## Author Contributions

SK SJW conceived, designed and performed the experiments; SK RS OG AN YS RC TK analyzed the data; SK TK contributed reagents/materials/analysis tools; SK RC TK wrote the paper. All authors reviewed the manuscript.

## References

1. Cooper, G. M. The Central Role of Enzymes as Biological Catalysts in The Cell: A Molecular Approach. 2nd edition. Sunderland (MA): (Sinauer Associates, 2000) NCBI Bookshelf ID: NBK9921 https://www.ncbi.nlm.nih.gov/books/NBK9921/

2. Berg, J. M., Tymoczko, J. L. & Stryer, L. Biochemistry. 5th edition. (NY) (Freeman W. H., 2002) NCBI Bookshelf ID: NBK21154 https://www.ncbi.nlm.nih.gov/books/NBK21154/

3. Anjem, A. & Imlay, J. A. Mononuclear iron enzymes are primary targets of hydrogen peroxide stress. J Biol Chem 287, 15544–15556 (2012).

4. Lodish, H. et al. Protein Structure and Function in Molecular Cell Biology 4th edition. (NY) (Freeman W. H., 2000) 963 NCBI Bookshelf ID: NBK21475 https://www.ncbi.nlm.nih.gov/books/NBK21475/

5. Bradbury, S. L. & Jakoby, W. B. Glycerol as an enzyme-stabilizing agent: effects on aldehyde dehydrogenase. Proc Natl Acad Sci USA 69, 2373–2376 (1972).

6. Carpenter, J. F., Martin, B., Loomis, S. H. & Crowe, J. H. Long-term preservation of dried phophofructokinase by sugars and sugar/zinc mixtures. Cryobiol 25, 372–376 (1988).

7. Argüelles, J. C. Physiological roles of trehalose in bacteria and yeasts: a comparative analysis. Arch Microbiol 174, 217–224 (2000).

8. Sakurai M. et al. Vitrification is essential for anhydrobiosis in an African chironomid, Polypedilum vanderplanki. Proc Natl Acad Sci USA 105, 5093–5098 (2008).

9. Crowe, J. H., Crowe, L. M., Carpenter, J. F. & Aurell Wistrom, C. Stabilization of dry phospholipid bilayers and proteins by sugars. Biochem J 242,1–10 (1987).

10. Green, J. L. & Angell, C. A. Phase relations and vitrification in saccharide-water solutions and the trehalose anomaly. J Phys Chem 93, 2880–2882 (1989).

11. Kreilgaard, L., Frokjaer, S., Flink, J. M., Randolph, T. W. & Carpenter, J. F. Effects of additives on the stability of Humicola lanuginosa lipase during freeze-drying and storage in the dried solid. J Pharm Sci 88, 281–290 (1999).

12. O’Fágáin, C. Storage and lyophilisation of pure proteins. Methods Mol Biol 681, 179–202 (2011).

13. Patel, S. M. & Pikal, M. J. Emerging Freeze-Drying Process Development and Scale-up Issues. AAPS PharmSciTech 12, 372–378 (2011).

14. Sales, K., Brandt, W., Rumbak, E. & Lindsey, G. The LEA-like protein HSP 12 in Saccharomyces cerevisiae has a plasma membrane location and protects membranes against desiccation and ethanol-induced stress. Biochim Biophys Acta 1463, 267–278 (2000).

15. Kikawada T. et al, Dehydration-induced expression of LEA proteins in an anhydrobiotic chironomid. Biochem Biophys Res Commun 348, 56–61 (2006).

16. Warner, A. H., Chakrabortee, S., Tunnacliffe, A. & Clegg, J. S. Complexity of the heat-soluble LEA proteome in Artemia species. Comp Biochem Physiol Part D Genomics Proteomics 7, 260–267 (2012).

17. Yamaguchi, A. et al, Two novel heat-soluble protein families abundantly expressed in an anhydrobiotic tardigrade. PLoS One 7, e44209 (2012).

18. Hinton, H. E. A new chironomid from Africa, the larva of which can be dehydrated without injury. J Zool 121, 371–380 (1951).

19. Watanabe, M., Kikawada, T. & Okuda, T. Increase of internal ion concentration triggers trehalose synthesis associated with cryptobiosis in larvae of Polypedilum vanderplanki. J Exp Biol 206, 2281–2286 (2003).

20. Mitsumasu, K. et al, Enzymatic control of anhydrobiosis-related accumulation of trehalose in the sleeping chironomid, Poiypediium vanderpianki. FEBS J 277, 4215–4228 (2010).

21. Hatanaka, R. et al, An abundant LEA protein in the anhydrobiotic midge, PvLEA4, acts as a molecular shield by limiting growth of aggregating protein particles. Insect Biochem Mol Biol 43, 1055–1067 (2013).

22. Gusev, O. et al. Anhydrobiosis-associated nuclear DNA damage and repair in the sleeping chironomid: linkage with radioresistance. PloS One 5, e14008 (2010).

23. Cornette, R. & Kikawada, T. The Induction of Anhydrobiosis in the Sleeping Chironomid: Current Status of Our Knowledge. IUBMB Life 63, 419–429 (2011).

24. Watanabe, K., Imanishi, S., Akiduki, G., Cornette, R. & Okuda, T. Air-dried cells from the anhydrobiotic insect, Polypedilum vanderplanki, can survive long term preservation at room temperature and retain proliferation potential after rehydration. Cryobiol 73, 93–98 (2016).

25. Sogame, Y. et al, Establishment of gene transfer and gene silencing methods in a desiccation-tolerant cell line, Pvll. Extremophiles 21, 65–72 (2017).

26. Tsien, R. Y. The green fluorescent protein. Annu Rev Biochem 67, 509–544 (1998).

27. Tapia, H. & Koshland, D. E. Trehalose is a versatile and long-lived chaperone for desiccation tolerance. Curr Biol 24, 2758–2766 (2014).

28. Borovikova, D., Teparić, R., Mrša, V. & Rapoport, A. Anhydrobiosis in yeast: cell wall mannoproteins are important for yeast Saccharomyces cerevisiae resistance to dehydration. Yeast 33, 347–53 (2016).

29. Kikuta, S., Hou, B. H., Sato, R., Frommer, W. B. & Kikawada, T. FRET sensor-based quantification of intracellular trehalose in mammalian cells. Biosci Biotechnol Biochem 80, 162–165 (2015).

30. Gstraunthaler, G. Alternatives to the use of fetal bovine serum: serum-free cell culture. ALTEX 20, 275–281 (2003).

31. Singh, J. et al, Transcriptional response of Saccharomyces cerevisiae to desiccation and rehydration. Appl Environ Microbiol 71, 8752–8763 (2005).

32. Gusev, O. et al, Comparative genome sequencing reveals genomic signature of extreme desiccation tolerance in the anhydrobiotic midge. Nat Commun 5, 4784 (2014). doi: 10.1038/ncomms5784.

33. Nakahara, Y. et al, Cells from an anhydrobiotic chironomid survive almost complete desiccation. Cryobiol 60, 138–146 (2010).

34. Hatanaka, R. et al, Diversity of the expression profiles of late embryogenesis abundant (LEA) protein encoding genes in the anhydrobiotic midge Polypedilum vanderplanki. Planta 242, 451–459 (2015).

35. Vancha, A. R. et al, Use of polyethyleneimine polymer in cell culture as attachment factor and lipofection enhancer. BMC Biotechnol 4, 23 (2004).

36. Rice, R. H. & Green, H. Relation of protein synthesis and transglutaminase activity to formation of the cross-linked envelope during terminal differentiation of the cultured human epidermal keratinocyte. J Cell Biol 76, 705–711 (1978).

37. Kikuta, S. et al, Characterization of a ligand-gated cation channel based on an inositol receptor in the silkworm, Bombyx mori. Insect Biochem Mol Biol 74, 12–20 (2016).

